# Gene Embedding Hub: a standardized resource for human gene representations

**DOI:** 10.64898/2026.10.29.762818

**Authors:** Daniel Tabares, Michael Bohl, Valentina Boeva

## Abstract

Gene embeddings are vector representations of gene identity that provide machine-learning models with information from diverse biological sources. However, their use is hindered by a lack of standardization: inconsistent identifiers, formats, and preprocessing conventions. In this work, we present a unified resource of 67 human gene-embedding sets derived from protein, DNA and RNA sequences, biomedical text, bulk expression data, single-cell expression models, functional data, biological networks, knowledge graphs, regulatory genomics and multimodal models, covering up to 20,097 GENCODE-annotated protein-coding genes. The collection includes both published precomputed embeddings and representations generated or extracted from publicly available models and biological data. Each embedding set is organized in a folder containing the embedding matrix, the Ensembl gene identifiers, machine-readable metadata, and processing documentation. We validate the resource by measuring global and local similarity, cross-embedding consensus, and cross-embedding predictability. The collection is designed to allow community-contributed embeddings that meet predefined technical and documentation requirements, enables comparison of gene representations across modalities and provides standardized embeddings for training and downstream applications.

## 1 Background & Summary

Genes can be described through their nucleotide and protein sequences, expression patterns, molecular interactions and accumulated functional knowledge. Gene embeddings translate information from these heterogeneous sources into numerical vectors that can be used in computational analyses and predictive models. By encoding relationships between genes, these embeddings can be useful for tasks including functional prediction [16], single-cell analysis [44, 52] and prediction of transcriptional responses to genetic perturbations [53]. Their role extends beyond reducing the dimensionality of biological data: when incorporated into downstream models, gene embeddings can introduce prior information that is absent from the measurements collected in an individual experiment. Protein-sequence embeddings are able to capture structural and evolutionary constraints [37], regulatory-sequence models include features linked to gene regulation [47], and expression-derived representations show the relationships that have been learned from cellular observations [44, 52]. Network-based methods provide a summary of molecular connectivity [36], while text-derived embeddings record associations found in descriptions of genes and their functions [39]. Since these different sources offer various views of gene biology, having a diverse set of them is useful for a wide range of applications with differing information needs.

Previous benchmarking studies have evaluated gene embeddings and released reusable collections. The Protein RepresentatiOn BEnchmark (PROBE) studied protein representations for functional prediction tasks [15], while later research compared gene embeddings obtained from network, text, sequence, and expression data [16]. Broad gene- property benchmarks have also allowed comparisons among different model families by means of common prediction tasks [54]. Their results show that performance varies according to the biological properties being assessed and the sources of information used for the embeddings [16, 54], thus underscoring the importance of developing a gene embedding resource that can be reused.

Despite recent progress, using and comparing embeddings from multiple sources still requires substantial harmonization. Embeddings are distributed across separate repositories and model releases, use different identifier conventions, and cover varying gene sets. Some are available as precomputed vectors, while others must be extracted from model checkpoints or generated from biological data. Additional challenges include differences in sequence selection, aggregation, and preprocessing, all of which affect the interpretation of gene-level vectors. Therefore, assembling a usable resource requires harmonizing these representations and documenting the processing decisions.

Here, we present the Gene Embedding Hub, a standardized collection of human protein-coding gene embeddings derived from protein, DNA and RNA sequences, biomedical text, bulk expression data, single-cell expression models, functional data, biological networks, knowledge graphs, regulatory genomics and multimodal models. The resource brings together published embeddings, others extracted from pretrained models and gene-level profiles constructed from biological data. A common gene-identifier framework and consistent organization support access and comparison, while accompanying documentation records the origin and processing of each embedding. The collection is publicly accessible at https://huggingface.co/spaces/BoevaLab/Gene-Embedding-Hub and is designed to accommodate additional community contributions. The Hub provides a reusable basis for benchmarking gene embeddings, evaluating alternative sources of biological information in predictive models, and studying different combinations across modalities.

## 2 Methods

### 2.1 Reference gene table construction and identifier harmonization

To keep gene identities consistent across the source embeddings, we built a master reference gene table that defines the reference set of human protein-coding genes considered in this study and establishes a link between each Ensembl gene identifier and the corresponding alternative gene, transcript, and protein identifiers. We prepared the table using protein-coding genes from the GENCODE release 49 annotation on the GRCh38 assembly [1, 2] and kept only those records with a GTF gene_type attribute of protein_coding. The Ensembl gene identifiers were converted to their stable forms by deleting the version suffixes, while the original versioned identifiers were retained in a separate column of the reference table for provenance. For each gene, the table includes its gene symbol, genomic coordinates, strand, and the associated transcript and protein identifiers.

We selected one representative transcript and its associated protein identifier for each gene using a fixed hierarchical procedure based on GENCODE transcript annotations, transcript support level, and total exon length, as described in Supplementary Methods. The selected transcript and protein identifiers were stored in the master reference table in both stable and versioned forms for use whenever a single transcript, coding sequence, or protein reference was required.

Stable Ensembl gene identifiers from the GENCODE-derived master reference table were submitted to MyGene.info [6, 7] using the human species filter to obtain Entrez Gene identifiers and UniProt accessions [10], preferring UniProtKB/Swiss-Prot over UniProtKB/TrEMBL. The resulting mappings were compared with the HGNC com- plete set and with an Ensembl BioMart query for protein-coding genes, Entrez Gene identifiers, and both UniProt accession types [8, 9]. The missing values in the main Entrez Gene field were only filled in if the combined HGNC and BioMart mappings provided a single unique identifier. When MyGene.info did not provide a UniProt mapping, all UniProt accessions reported for that gene by HGNC and BioMart were retained.

We extended identifier coverage by combining the HGNC annotations with the transcript and protein identifiers from Ensembl release 116 on GRCh38 and Ensembl release 87 on GRCh37 [4, 14]. HGNC records were matched to the reference table using Ensembl gene identifiers, approved gene symbols, Entrez Gene identifiers, and UniProt accessions, and were included in the reference table only when all identifiers that matched the reference table pointed to the same reference gene. Consolidated lookup fields were then set up separately for the approved, previous, and alias gene symbols as well as for the Ensembl gene names; for the Entrez Gene and UniProt identifiers; for the RefSeq and CCDS identifiers [11, 12]; for the stable and versioned Ensembl transcript and protein identifiers; and for the STRING-compatible protein identifiers [13]. Those HGNC records that did not link to any reference gene were omitted. The final lookup fields connect the source-specific identifiers to the stable, unversioned Ensembl gene identifiers.

### 2.2 Embedding selection, processing, and standardization

We compiled 67 human gene-embedding sets by either downloading public precomputed embedding matrices from previous studies or generating them ourselves from public data or released models. Potential resources were primarily identified in research papers, published benchmarking collections [15, 16], and public biological datasets, and were included only if they offered fixed-length numerical vectors for each represented human gene or provided enough data to generate these vectors. Embeddings created using different algorithms, training sources, or model checkpoints were treated as separate sets. For each embedding set, we documented its modality, gene coverage, dimensionality, acquisition route, and processing procedure (Supplementary Table 1).

Precomputed embeddings were modified to harmonize identifiers and ensure technical compatibility. We sourced the matrices and their identifiers from repositories linked to original publications, preserving both their original dimensionality and numerical values. Source identifiers were standardized by type and matched to the corresponding entries in the master gene table. Source rows without a match or mapped to multiple reference genes were excluded, whereas vectors from multiple source rows that mapped uniquely to the same reference gene were averaged following the application of any source-specific mapping priorities. Identifier harmonization outcomes were recorded for each precomputed embedding set (Supplementary Table 2).

Locally generated embeddings were obtained by applying released models to gene-associated inputs, constructing representations from structured biological data, or extracting fixed gene representations from public checkpoints. Inputs included RNA transcript sequences, DNA coding sequences, protein sequences, and GRCh38 genomic regions surrounding transcription start sites or splice sites [2]. Other inputs included gene and protein descriptions, matrices of expression or functional data, and network or knowledge-graph data. When models produced token- or residue- level embeddings, we averaged the vectors from the selected model layer, excluding padding and special tokens. Sequences exceeding model context limits were processed in non-overlapping chunks and combined using token-count or sequence-length weights. Source-specific transformations, dimensionality reduction, and model settings are detailed in Supplementary Table 1 and the per-embedding README files. Locally generated embeddings were aligned with the master gene table. When embedding vectors were generated for genes selected from the master table, we used their unversioned Ensembl gene identifiers during input selection and kept them in the final output. When they were instead indexed with external identifiers, we applied the same mapping and aggregation rules as for precomputed matrices. Genes without valid representations were excluded.

Identifier harmonization produced 67 standardized embedding sets, each indexed by stable Ensembl gene identifiers and containing one vector per retained gene (Fig. 1). Gene coverage and dimensionality vary across sets, reflecting differences in the source data and representation methods.

**Figure 1.**
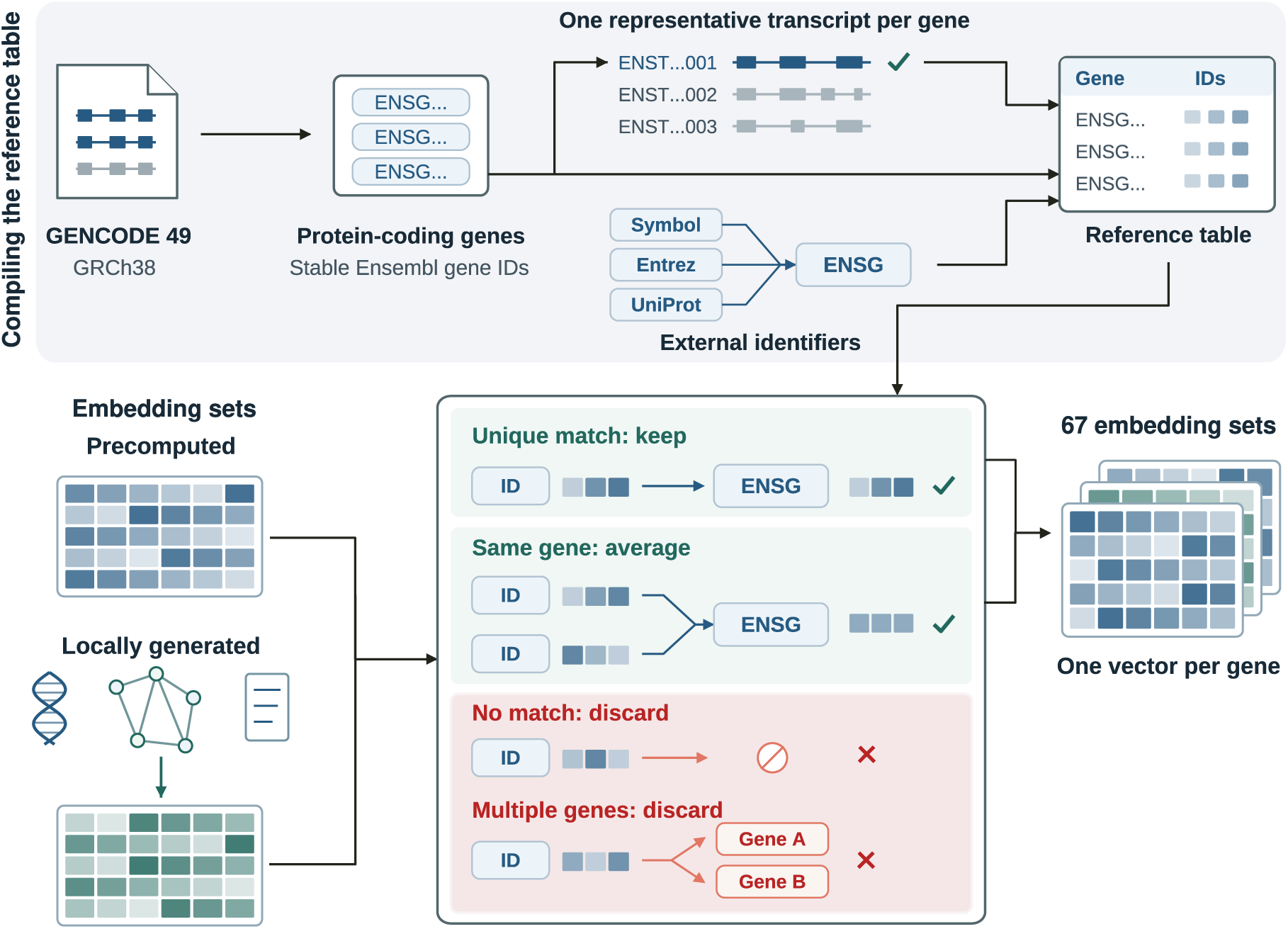
Overview of the reference gene table construction and gene embedding standardization. Human protein-coding genes from GENCODE v49 (GRCh38) were linked to representative transcripts and external identifiers. Precomputed and locally generated embeddings were mapped to stable Ensembl gene identifiers. Uniquely mapped vectors were retained, multiple vectors mapping to the same gene were averaged, and unmatched or ambiguously mapped vectors were discarded. This yielded 67 standardized embedding sets, each containing one vector per retained gene.

### 2.3 Computational implementation

Data acquisition, model inference, identifier harmonization, matrix conversion, and quality-control procedures were implemented in Python, using NumPy [17] and pandas [18] for numerical and tabular data processing. PyTorch [19] was used to implement pretrained model inference and checkpoint extraction, either directly, via the Hugging Face Transformers library [20], or through specific model implementations. Due to incompatible software dependencies among several original models, the corresponding pipelines were run in separate virtual or Conda environments. Nearly all tasks were performed on CPUs (Intel Core i9-9900K processor or AMD EPYC 7H12) with up to 64 GB of RAM. Neural network models were run on NVIDIA GeForce RTX 2080 Ti GPUs with 11 GB of VRAM. Details about processing parameters, including the representation layer, pooling or aggregation procedure, sequence-chunking and batch settings, numerical precision, and random seed, when applicable, are provided in the embedding documentation. The resource interface loads a generated JSON catalog and was built as a static Hugging Face Space using HTML, CSS, and JavaScript.

## 3 Data Records

### 3.1 Resource structure

The **Gene Embedding Hub** dataset repository is available on Hugging Face: https://huggingface.co/datasets/BoevaLab/Gene-Embedding-Hub-dataset, along with a space that enables users to browse the different embeddings and obtain the related files: https://huggingface.co/spaces/BoevaLab/Gene-Embedding-Hub.

Each downloadable embedding set is distributed in a separate directory containing four files: embeddings.npz, genes.tsv, metadata.json, and README.md.

The embeddings.npz file contains the numerical representation matrix as a compressed NumPy archive. The matrix is stored under the key embeddings, with rows corresponding to genes and columns varying based on the number of features in the representation.

The genes.tsv file contains a single ensembl_gene_id column containing stable, unversioned Ensembl gene identifiers. The order of these gene identifiers matches the order of the rows in the embedding matrix and therefore correctly maps genes to vectors. Gene coverage varies between different embeddings, and only genes that have a valid representation are included in each file. Consequently, the absence of a gene from a given genes.tsv file denotes the absence of a representation for that gene and should not be interpreted as a zero-valued vector.

The metadata.json file provides machine-readable documentation for each embedding. It describes the number of genes, embedding dimensionality, modality, acquisition route, and coverage of the master gene reference set. Additional representation-specific fields may be included depending on the embedding.

The README.md file describes the inputs and procedure followed to obtain each embedding. It also details processing and transformation decisions, model checkpoints or data releases, procedures for mapping identifiers, quality control information, and links to the source resources and related publications.

Since different representations do not necessarily include the same genes, comparisons or combinations of the embedding matrices should be carried out using the Ensembl gene identifiers found in their respective genes.tsv files. Their gene lists can be intersected and the corresponding matrices rearranged so that matched gene rows are obtained.

### 3.2 Community contributions

The Gene Embedding Hub supports community contributions of additional human gene embeddings through a standardized workflow. Contributors have to upload a directory named with the embedding_id to the Hugging Face contributions Dataset and submit it as a pull request. Each submission must contain four files: embeddings.npz, genes.tsv, README.md, and submission.yaml.

The first three files must meet the matrix, gene identifier, and documentation requirements described earlier. The submission.yaml file is a form that must be completed before submitting the package and includes scientific context, provenance, licensing information, and contributor statements. The template can be directly downloaded from the Gene Embedding Hub. Since the contributed files must use Ensembl gene identifiers, a gene-mapping tutorial is included in the resource. Optional license files may also be included.

Open pull requests are handled using a GitHub Actions pipeline that verifies the technical requirements detailed in the Technical Validation section. If the submitted package successfully passes all the checks, it is automatically merged into the public dataset under data/community_contributions/<embedding_id>. A metadata.json file is automatically generated following the same structure as the metadata.json files in the core dataset and saved alongside the other files in the same folder. The Space catalog is also updated to show the contributed embedding in the Gene Embedding Hub, and the pull request is then closed. If any validation check fails, the contributor receives a diagnostic report through the pull request with the failed checks and the public dataset stays unchanged.

## 4 Data overview

The core dataset contains 67 human gene embeddings across 11 modality categories based on the principal type of biological information each embedding encodes (Fig. 2a). These categories cover embeddings derived from protein, DNA, and RNA sequences, regulatory genomic context, bulk expression measurements and single-cell expression models, heterogeneous biomedical knowledge graphs (encoding relations between genes, proteins, diseases, drugs, phenotypes, pathways, and anatomical structures), molecular interaction networks, functional annotations and assays, biomedical text, and multimodal combinations of biological information (Supplementary Table 3). Individual embeddings cover between 13,420 and 20,097 genes from the set of 20,097 considered genes, and range from 20 to 20,421 dimensions (Fig. 2b).

**Figure 2.**
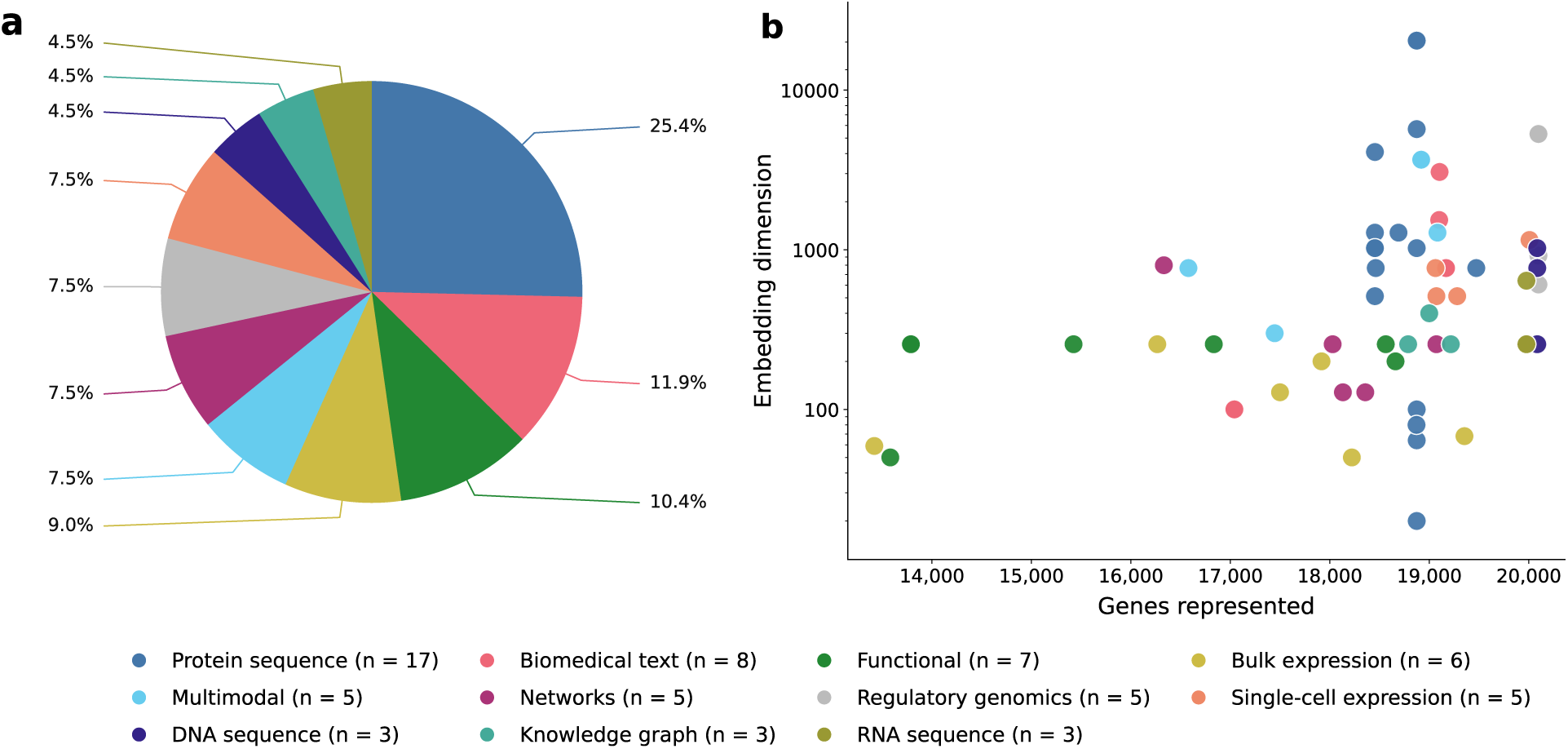
Statistical summaries of the core Gene Embedding Hub collection. **a.** Distribution of the 67 core embeddings across the 11 modalities defined in Supplementary Table 3. **b.** Gene coverage and dimensionality of the individual embeddings.

## 5 Technical Validation

### 5.1 Quality control of the core embedding collection

Each embedding underwent structural and numerical quality checks to ensure the integrity of the files. The number of rows in the final embedding matrix had to match the number of gene identifiers in the corresponding genes.tsv file, and the matrix’s dimensionality had to match the expected dimensionality of the downloaded source or the generation method. The genes.tsv file was checked for duplicated Ensembl gene identifiers and the numerical matrix was screened for NaN and infinite values or rows with Euclidean norm equal to zero, none of which were permitted.

### 5.2 Automated validation of community contributions

A scheduled GitHub Actions workflow automatically detects new pull requests on Hugging Face and uses trusted code from the default branch of the contributions repository. Each pull request must add a single embedding directory containing the four files specified in Data Records, with a directory name matching the submitted embedding_id.

The submission.yaml template must be downloaded from the resource and filled out with the embedding’s name, category, modality, and description. Required provenance fields, including the identifier type prior to harmonization and the mapping procedure, must also be completed. Explicit permission for redistribution is mandatory. The accuracy of the submitted information, rights regarding redistribution, absence of sensitive human and patient-level data, review of original source terms, and acceptance of automated checks must be confirmed in the file. Providing a source-license identifier is optional. However, if included, it must be listed on the automatic allowlist, which comprises Apache-2.0, BSD-3-Clause, CC-BY-4.0, CC0-1.0, MIT, or ODC-BY-1.0.

The genes.tsv file must include an ensembl_gene_id column with unique, non-empty values matching the stable Ensembl gene pattern: ENSG followed by 11 digits. These genes must represent at least 5% of the overall reference gene set, and no more than 2% are allowed to be outside of it.

The uncompressed embeddings.npz file should be less than 5 GB, unencrypted, and contain exactly one array named embeddings. This array must be loaded directly without object deserialization, be two-dimensional, use float32, have between 2 and 50,000 features, and have the same number of rows as genes.tsv. All matrix values must be finite, and no vector may have zero norm. Exact duplicate vectors and constant feature columns are not allowed. The README.md file must contain at least 200 bytes of documentation.

To prevent duplicate embedding archives in the resource, the proposed embedding_id is verified against the community registry and the core dataset catalog. The SHA-256 checksum of the embeddings.npz file is also compared with the checksums of the community contributions already registered, and any archive that has the same checksum is rejected.

### 5.3 Global and local concordance between embedding spaces

To assess whether matrix processing and identifier harmonization preserved meaningful relationships among genes, we compared the embedding spaces using two complementary metrics. We expected embeddings derived from related biological inputs or modeling strategies to preserve similar global gene–gene geometry and local gene neighborhoods [21, 22].

In order to use the same set of genes in every pairwise comparison, we determined the largest subset of embeddings whose strict intersection included at least 14,000 genes using an exact mixed-integer optimization. This optimization minimized the number of embeddings excluded while satisfying the threshold and, when several feasible subsets had the same size, it selected the one with the highest number of genes in their common intersection. This procedure retained 56 of the 67 embeddings and produced a common intersection of 14,039 genes, yielding 1,540 unordered pairs of embeddings. Since the same gene set was used for all pairs, differences in similarity could not be due to varying gene coverage.

For each pair, global concordance was quantified using unbiased linear centered kernel alignment (CKA) to the row-normalized gene vectors [21, 23]. CKA measures the agreement between the overall geometries produced by two representations, even if they have different dimensionalities. Local concordance was evaluated by creating exact cosine- similarity gene-neighborhood graphs and calculating the tie-aware overlap between the *k* = 100 [22] nearest-neighbor groups derived from each embedding (around 0.7% of the entire 14,039-gene set). Boundary ties were given fractional weights to prevent tied similarities from causing arbitrary neighbor selections. For each metric, a negative-control baseline was obtained from five deterministic gene-identity shuffles for each pair of embeddings. The scores were calculated as (observed similarity *−* mean shuffled similarity)/(1 *−* mean shuffled similarity) and were used to estimate and eliminate any chance overlap.

The CKA and neighborhood-overlap matrices showed recurring broad blocks of related embeddings, while also displaying metric-specific differences (Supplementary Fig. 1). We calculated Spearman’s correlation to quantify the agreement between the selected calibrated CKA and the calibrated *k* = 100 local-overlap view, and the result was tested using a quadratic assignment procedure (QAP) matrix-correlation test [24] with 9,999 separate permutations of both the rows and columns of the embedding labels. This permutation approach preserves the relationships between similarity pairs sharing the same embedding. The two views were positively correlated (Spearman’s *ρ* = 0.814; Holm-adjusted two-sided empirical *P* = 1.0 *×* 10*^−^*^4^). Robustness was tested by eliminating each embedding one at a time, and in all 56 of the leave-one-embedding-out analyses the correlation was still positive (median *ρ* = 0.815, ranging from 0.793 to 0.829). Together, the two views shared a consistent common signal while remaining sufficiently complementary to justify their combination.

Both similarity measures were placed on a common scale by independently converting their values across the 1,540 pairs to tie-aware percentile ranks. These values were then averaged with equal weights, yielding a consensus score that represented relative affinity. A weighted graph was constructed by connecting each embedding to its ten highest-scoring consensus neighbors, forming the symmetrized union of these directed neighborhoods, and using the consensus affinities as the edge weights. Communities were identified using deterministic greedy weighted-modularity optimization at resolution 1 [25]. Modality labels were excluded from graph construction, community detection, and layout, and were incorporated only after these steps for interpretative purposes (Fig. 3a).

**Figure 3.**
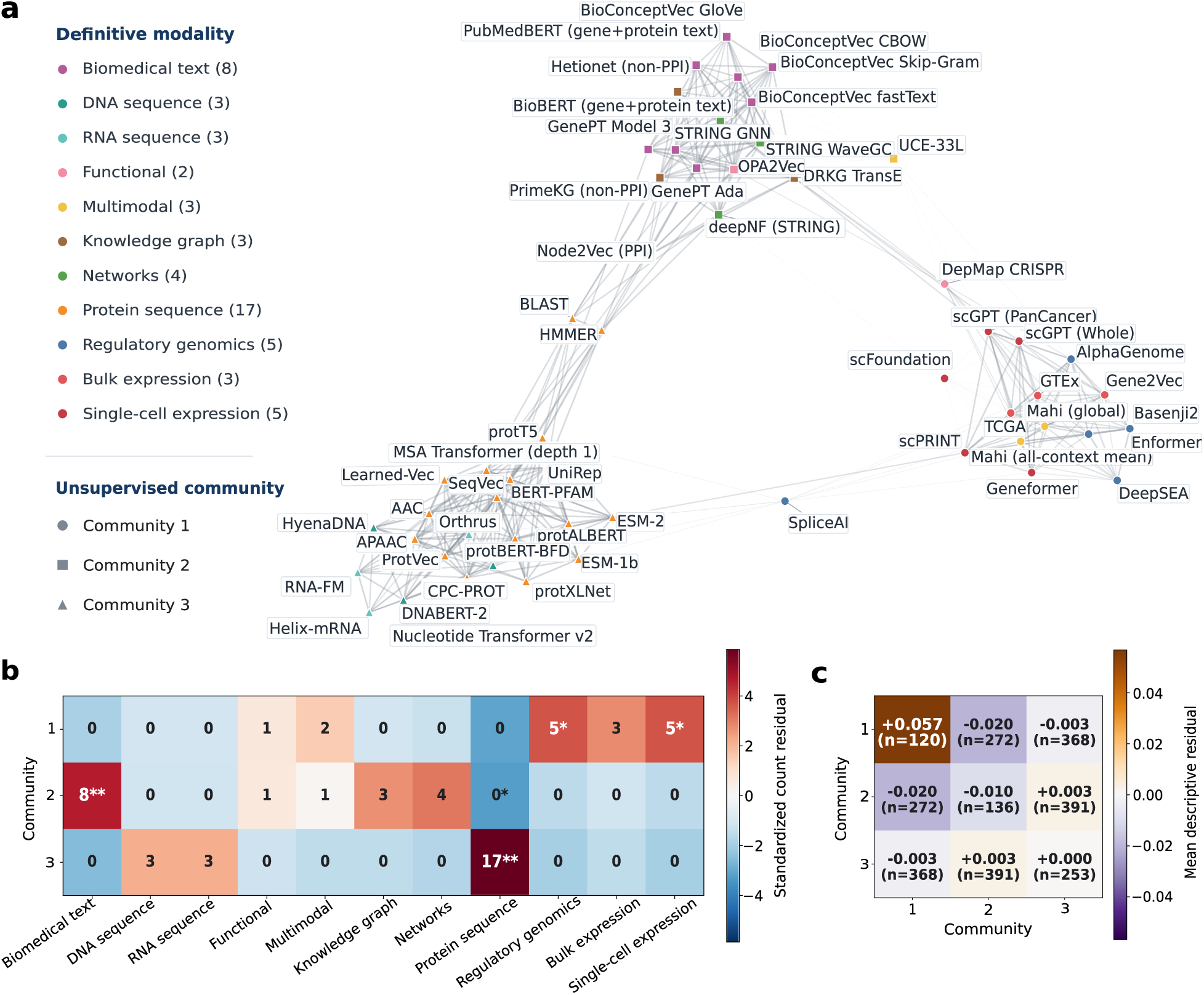
Consensus organization of gene-embedding spaces. **a.** Weighted, symmetrized 10-nearest-neighbor graph of 56 embeddings across 14,039 common genes. Edge weights reflect the equal-rank consensus of calibrated linear CKA and local- neighborhood overlap at *k* = 100. **b.** Community-by-modality composition. Colors show standardized deviations from the counts expected given the fixed community and modality sizes, with positive and negative values indicating enrichment and depletion, respectively. Cell labels indicate the observed number of embeddings, and asterisks denote community–modality associations identified by node-label permutation testing after Holm correction (* adjusted *P* < 0.05; ** adjusted *P* < 0.01). **c.** Mean modality- adjusted consensus affinity within and between communities. For each embedding pair, the mean consensus affinity of all other pairs with the same combination of modalities was subtracted from its consensus affinity. Positive and negative values indicate stronger and weaker similarity, respectively, than the corresponding modality-matched baseline. Cell labels show the mean residual affinity and the number of embedding pairs.

The resulting graph separated the 56 embeddings into three communities containing 16, 17, and 23 embeddings. Their composition showed agreement with the independently assigned modalities. Community 1 contained all five regulatory- genomics, all five single-cell expression and all three bulk expression embeddings, together with DepMap CRISPR and the two Mahi representations. Community 2 contained all biomedical-text, network, and knowledge-graph embeddings, together with OPA2Vec and UCE-33L. Community 3 contained all 17 protein-sequence, all three DNA-sequence and all three RNA-sequence embeddings. Thus, nine of the eleven modality categories were entirely confined to a single community. Fixed-count node-label permutations confirmed significant enrichment of regulatory-genomics and single-cell expression embeddings in Community 1 (adjusted *P* = 0.0493 and 0.0155, respectively), biomedical-text embeddings in Community 2, and protein-sequence embeddings in Community 3 after Holm correction [26] across the 33 community-by-modality combinations (both adjusted *P* = 0.0033, Fig. 3b). Protein-sequence embeddings were also significantly depleted from Community 2 (adjusted *P* = 0.0270). The complete confinement of several smaller modalities, including DNA sequence, RNA sequence, networks and knowledge graphs, did not always reach cell-wise significance after multiple-testing correction (all adjusted *P ≥* 0.1404) because these categories contained few embeddings.

The separation of the functional and multimodal embeddings into different communities is biologically plausible because these labels encompass representations constructed from substantially different information sources. Within the Functional category, DepMap CRISPR was obtained from gene-effect profiles measured across cancer cell lines and therefore captures context-dependent cellular responses [27], explaining its placement alongside bulk expression, single-cell expression, and regulatory-genomics embeddings in Community 1. In contrast, OPA2Vec encodes genes using Gene Ontology annotations, formal ontology structure, and textual metadata encoded using a model pretrained on PubMed [28]. Its association with biomedical-text, knowledge-graph, and network representations in Community 2 is therefore expected. Similarly, the two Mahi embeddings summarize a model integrating tissue-specific networks with chromatin accessibility, transcription-factor binding, histone modifications, and protein-derived features across multiple cellular contexts, resulting in strong regulatory and context-dependent content [29]. UCE-33L instead combines single-cell transcriptomic training with protein-sequence-derived gene representations [30]. As a result, the geometries induced by the Mahi embeddings and UCE-33L are not expected to be identical.

Finally, we examined whether the detected communities showed structural features beyond those expected from their modality composition. For each embedding pair, the mean consensus similarity of the exact modality-pair block was deducted by using a leave-one-pair-out baseline. Community 1 retained a positive mean residual affinity of +0.057, while Communities 2 and 3 showed values of *−*0.010 and around 0.000, respectively (Fig. 3c). The mean residual affinities between the communities were small or negative (*−*0.020, *−*0.003, and +0.003). These results indicate additional cross-modality coherence in Community 1, whereas the structure of Communities 2 and 3 is mainly due to their strongly aligned modality composition.

### 5.4 Directional predictability between embedding spaces

To further assess the embedding collection, we performed a directional reconstruction analysis to determine whether the information represented in one space can be recovered from another. We expected embeddings derived from closely related inputs to be mutually predictable and embeddings integrating broader biological information to reconstruct more specialized representations better than the reverse.

All pairs of 67 embeddings were evaluated in both directions, resulting in 4,422 source-to-target prediction tasks, with genes split into 70% training, 10% validation, and 20% test sets. For each direction, source vectors were standardized using only the training genes and subsequently used to predict a principal component analysis (PCA) representation of the target embedding. The PCA of the target embedding was also fitted using only the training data, selecting principal components to retain at least 90% of the target variance, up to a maximum of 1,024 components. Multilayer perceptrons with one to four hidden layers were trained for each direction, with the final model depth selected based on the *R*^2^ value on the validation set. Performance was reported as the full-target test *R*^2^, calculated relative to the total variance of the original target embedding, thus accounting for variance discarded during PCA. This procedure prevented embeddings that were easier to compress from receiving an artificial advantage. A value of zero indicates that the target training mean was predicted. Additional details of the MLP architecture and optimization procedure are provided in the Supplementary Technical Validation section.

The resulting source-to-target matrix was highly directional (Fig. 4a). As an internal positive control, the two Mahi representations (all-context mean and global) were highly mutually predictable and both achieved the highest median outgoing *R*^2^ values within the entire panel, which aligns with the expectation that embeddings integrating information from multiple biological contexts should recover more specific embeddings more easily. The network- fusion embedding deepNF [31] ranked next, while other broad network and knowledge-graph representations, including PrimeKG, STRING WaveGC, DRKG, Hetionet, and Mashup [13, 32, 33, 34, 35, 36], also exhibited substantial outgoing predictability. The positions of these representations are biologically reasonable since interaction networks and knowledge graphs aggregate relationships derived from several underlying evidence types.

**Figure 4.**
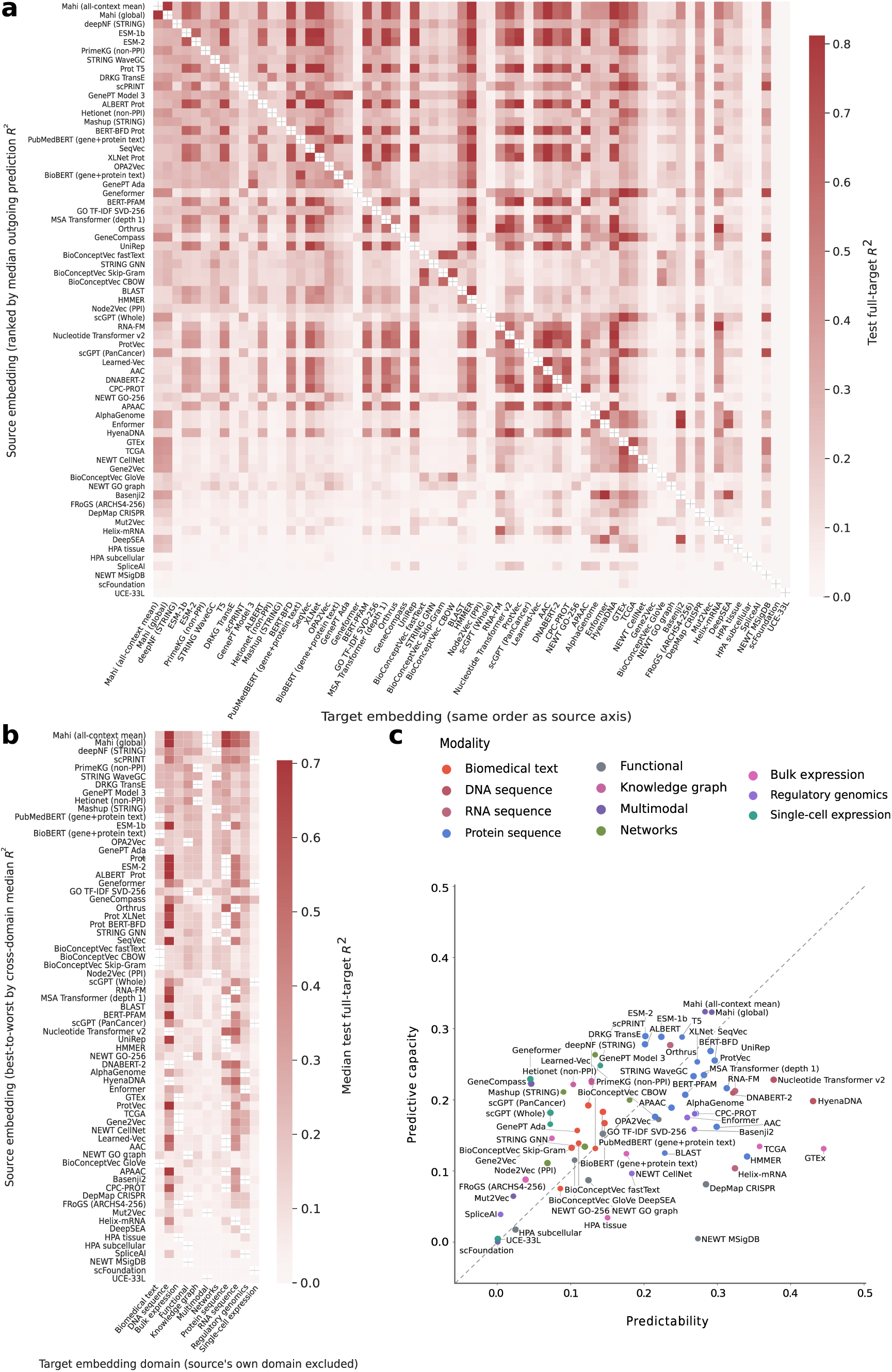
Directional cross-embedding reconstruction. **a.** Source-to-target reconstruction performance across all 67 gene embeddings. Each cell shows the *R*^2^ obtained by predicting the target embedding from the source embedding using the validation- selected multilayer perceptron; rows correspond to source embeddings and columns to targets, so asymmetry reflects directional predictability. **b.** Cross-domain reconstruction *R*^2^ after excluding targets belonging to the source embedding’s own modality. For each source–target-modality combination, values represent the median *R*^2^ across embeddings in the target modality. **c.** Predictive capacity versus predictability for each embedding, calculated by taking the median *R*^2^ for each target modality and averaging these values across modalities, using outgoing predictions for predictive capacity and incoming predictions for predictability.

A second positive control was provided by the protein-sequence embeddings. ESM-1b and ESM-2 were both among the five highest-ranked sources, with strong prediction between the two related protein language-model spaces [16, 37]. protT5, protALBERT, protBERT-BFD, SeqVec, protXLNet, BERT-PFAM, the MSA Transformer and UniRep also displayed broad outgoing signal [15, 16, 38]. In contrast, simpler sequence summaries such as amino-acid composition (AAC), pseudo-amino-acid composition (APAAC), ProtVec, and Learned-Vec achieve lower rankings [15, 16]. This was again expected, since embeddings obtained from large protein-sequence corpora capture a wider range of sequence and evolutionary structure compared to fixed-composition or short-context encodings.

Similar patterns appear within other modalities. GenePT (Model 3), PubMedBERT and BioBERT gene-and-protein-text embeddings were more broadly predictive than most of the lower-dimensional BioConceptVec embeddings, consistent with their use of richer gene and protein descriptions [39, 40, 41, 42]. Among single-cell RNA-seq-derived embeddings, scPRINT and Geneformer generalized effectively to embeddings outside the single-cell expression modality, indicating that their learned gene representations contain co-expression or regulatory relationships also present in other biological data sources [43, 44, 45]. Within regulatory genomics, the broader AlphaGenome and Enformer track summaries ranked above the more narrowly defined Basenji2, DeepSEA and SpliceAI embeddings [46, 47, 48, 49, 50].

Modality provided more information than dimensionality, as modality-only and dimensionality-only models explained 44.9% and 15.2% of the variation in directional predictability, respectively. A quadratic assignment procedure regression of their joint association was significant (9,999 permutations; empirical *P* = 10*^−^*^4^), indicating that structured differences among embeddings were preserved after harmonization.

To evaluate cross-domain transfer directly, we summarized each source against target modalities while excluding targets from the source’s own modality (Fig. 4b). The two Mahi embeddings remained the strongest overall, while deepNF and several knowledge-graph embeddings also demonstrated broad cross-domain performance. Protein language models exhibited particularly strong recovery of DNA- and RNA-sequence representations. Additionally, GenePT Model 3 and gene-and-protein-text embeddings effectively transferred signal to functional, network, and sequence-derived targets. The persistence of these patterns after removal of same-modality targets indicates that recoverable structure extends across the assigned modality labels.

We next distinguished each embedding’s ability to predict other spaces from its tendency to be predicted by them (Fig. 4c). Predictive capacity and predictability for each embedding were determined by calculating the median *R*^2^ for each target modality and averaging these values across modalities. Embeddings above the diagonal contain less common or broadly transferable information, while embeddings below the diagonal are easier to reconstruct but offer less recoverable information, indicating greater redundancy or specialization.

ESM-2, ESM-1b, protT5, deepNF, scPRINT, Geneformer, and GeneCompass appeared above the diagonal, indicating that these models provide broad or distinct information. The two Mahi representations combined the highest predictive capacity with high, but lower, predictability. Conversely, GTEx and TCGA were among the most predictable targets but had substantially lower outgoing capacity. Their co-expression profiles therefore appear to capture biological variation that is represented across several other modalities, while contributing less information that can be reconstructed broadly outside expression-related spaces. HyenaDNA showed a similar but less pronounced pattern. DepMap CRISPR and NEWT MSigDB were also highly predictable by other embeddings in the set but offered lower predictive capacity, which aligns with their more limited focus on gene-dependency phenotypes and their respectively curated gene-set structure [27, 51].

Finally, directional reconstruction demonstrated strong concordance with the two symmetric measures used in the preceding analysis (Supplementary Fig. 2). Among all 2,211 embedding pairs, the bidirectional mean selected-MLP *R*^2^ exhibited a strong correlation with linear CKA (*ρ* = 0.764, *P* < 10*^−^*^300^) and nearest-neighbor overlap (*ρ* = 0.719, *P* < 10*^−^*^300^). Thus, pairs that shared global geometry or local gene neighborhoods were also easier to translate between. The improvement of the selected nonlinear model over the linear baseline (no hidden layers) was associated with CKA (*ρ* = 0.568, *P* = 2.4 *×* 10*^−^*^189^) and neighbor overlap (*ρ* = 0.628, *P* = 2.1 *×* 10*^−^*^242^), indicating that nonlinear gains were concentrated among biologically related spaces rather than occurring randomly.

Together, these results offer three complementary technical checks. Related embeddings were predictably recon- structable, information-rich sources enabled broad outgoing predictions across modalities, and pairwise scores aligned with independent measures of global and local concordance. Directional reconstruction therefore recovers expected relations within the gene representations, supporting both the biological and technical validity of the embedding collection.

## 6 Data availability

Gene Embedding Hub is available at https://huggingface.co/datasets/BoevaLab/Gene-Embedding-Hub-dataset, with a web interface at https://huggingface.co/spaces/BoevaLab/Gene-Embedding-Hub. Matrices are distributed only where the applicable terms permit redistribution. For entries without downloadable matrices, source links and, where permitted, scripts for acquisition or local generation and standardization are provided. Third-party data, software, and model weights remain subject to their original licenses, terms of use, and access requirements.

## 7 Code availability

Custom code for acquiring, generating, standardizing and checking the embeddings, as well as reproducing the technical validation, is available at https://github.com/DanielTabaresLopez/Gene-Embeddings. The necessary code to implement the Gene Embedding Hub web interface is available at https://github.com/DanielTabaresLopez/Gene-Embedding-Hub-contributions. The repository includes usage instructions, software dependencies and configuration details.

## 8 Author contributions

D.T. developed the computational framework, implemented the code, performed the analyses, assembled the resource, and wrote the original manuscript draft. M.B. supervised the work, provided guidance on methodology and implemen- tation, and contributed to manuscript revision and editing. V.B. conceived the project and contributed to manuscript review and editing. All authors read and approved the final manuscript.

## Supplementary material

### Supplementary Methods: Reference data sources

The reference files used to build the master gene table were downloaded from the GENCODE release 49 annotation on GRCh38 (https://ftp.ebi.ac.uk/pub/databases/gencode/Gencode_human/release_49/gencode.v49.annotation.gtf.gz), the HGNC complete set (https://www.genenames.org/download/statistics-and-files/ ), the Ensembl release 116 GTF on GRCh38 (https://ftp.ensembl.org/pub/release-116/gtf/homo_sapiens/Homo_sapiens.GRCh38.116.gtf.gz), and the Ensembl release 87 GTF on GRCh37 (https://ftp.ensembl.org/pub/grch37/release-87/gtf/homo_sapiens/Homo_sapiens.GRCh37.87.gtf.gz). The additional identifier mappings were obtained by querying the MyGene.info v3 API (https://docs.mygene.info/en/latest/doc/query_service.html) through its */v3/query* endpoint and Ensembl BioMart (https://www.ensembl.org/info/data/biomart/index.html) through its */biomart/martservice* endpoint. All the files were downloaded and all the API and database queries were performed on July 3, 2026.

### Supplementary Methods: Representative transcript and protein selection

We used a fixed hierarchical selection procedure to assign to each gene one representative transcript together with its associated protein identifier. If one or more transcripts had a GTF transcript_type attribute of protein_coding, selection was restricted to those transcripts. For the 351 genes without a transcript carrying that exact classification, all associated transcripts were considered. Subsequently, the retained transcripts were ranked hierarchically using three representative-transcript annotations recorded in the GTF. Transcripts carrying the MANE Select tag were assigned the highest priority [3]. Among the remaining transcripts, those carrying the Ensembl canonical tag were ranked next, followed by those with an APPRIS principal tag [4, 5]; transcripts carrying none of these three tags were ranked last. Within each annotation category, transcripts were further ranked using the GENCODE transcript_support_level attribute, which summarizes how well their full transcript structure is supported by mRNA and expressed sequence tag alignments. Lower ranks were prioritized, whereas missing, NA, or other non-numeric values were ranked last. Transcripts remaining tied within a category were then ranked by total exon length, calculated as the sum of their exon intervals, with longer transcripts preferred. For each of the 351 genes without transcripts with the protein_coding attribute, the chosen transcript had an annotated coding sequence and protein identifier.

### Supplementary Technical Validation: MLP architecture and training

For each ordered source-to-target embedding pair, the source matrix was standardized using the feature-wise mean and standard deviation calculated from the training genes only. Target representations were centered using their training-set mean and divided by a single training-set root-mean-square scale. Target dimensionality was reduced using PCA fitted exclusively to the training genes. The smallest number of components explaining 90% of the training variance was retained, with a minimum of 16 and a maximum of 1,024 components.

Nonlinear predictors contained between one and four hidden layers. Each hidden layer consisted of a linear trans- formation followed by layer normalization, a GELU activation and dropout with probability 0.1. Dropout was also applied to the input. The hidden width was selected separately for each source-to-target direction as the smallest value among 32, 64, 128, 256 and 512 that was at least as large as the greater of the source’s intrinsic dimensionality and the number of retained target components; the width was capped at 512. A linear model (no hidden layers) using the same preprocessing served as a linear baseline.

Models were optimized by minimizing mean-squared error with AdamW. Nonlinear models used an initial learning rate of 10*^−^*^3^, whereas the linear model used 3 *×* 10*^−^*^4^. Weight decay was 10*^−^*^4^, the batch size was 256 and the gradient norm was clipped at 1.0. Learning rates followed a cosine-annealing schedule over a maximum of 300 epochs. Training was stopped when validation *R*^2^ failed to improve by more than 10*^−^*^5^ for 20 consecutive epochs, and the parameters from the epoch with the highest validation score were restored. For each direction, the final nonlinear architecture was selected among depths one to four according to full-target validation *R*^2^, with ties resolved in favor of the shallower model. The held-out test set was not used for either early stopping or architecture selection.

The reported full-target score was calculated as 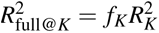 , where 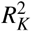 is the coefficient of determination in the retained *K*-component target subspace and *f_K_* is the fraction of the original target test variance represented by those components. This is equivalent to predicting omitted target directions using their training-set means. Scores were not truncated at zero.

## Supplementary Figures

**Supplementary Figure 1.**
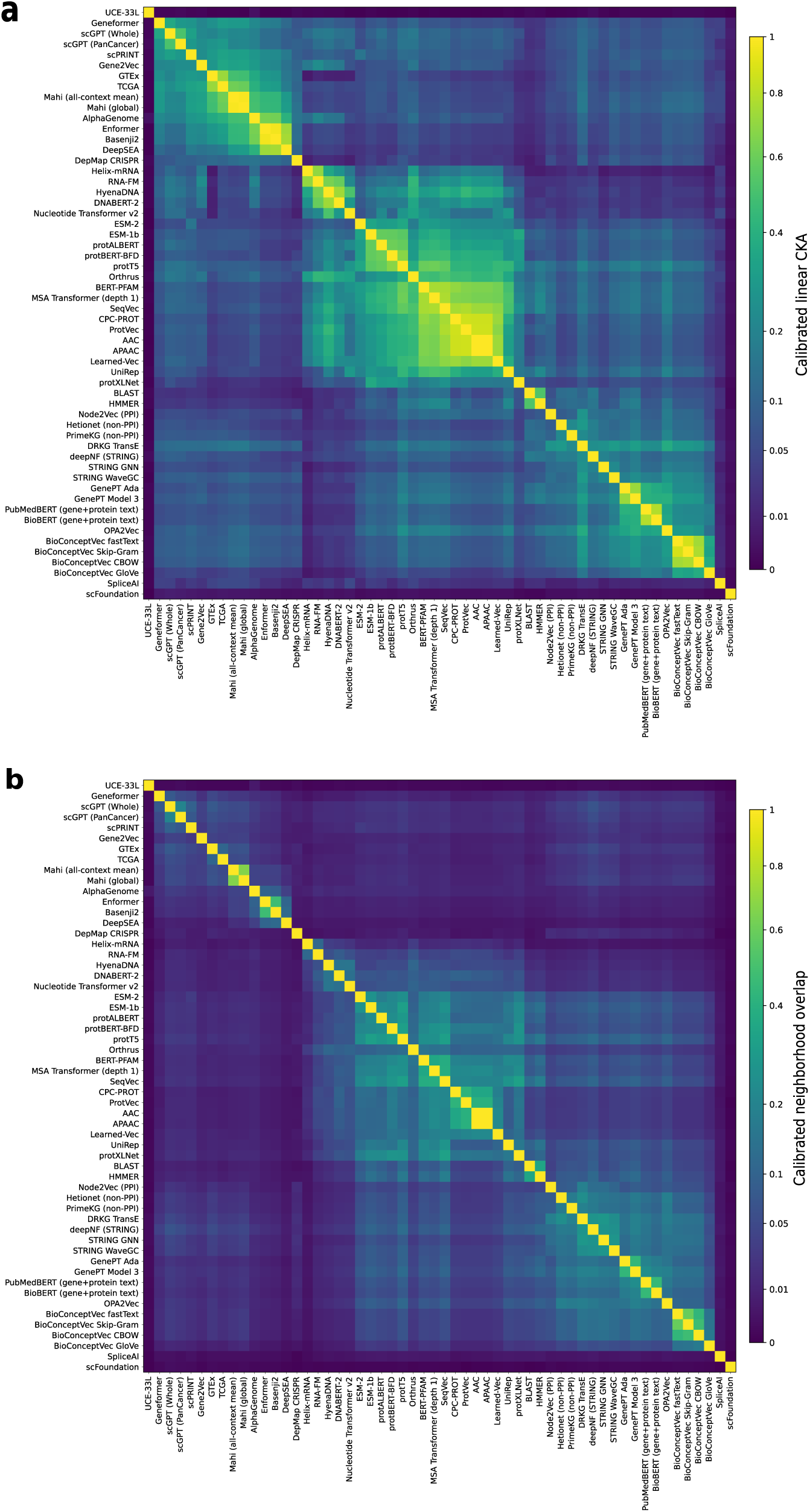
Global and local concordance among gene embeddings. **a.** Pairwise values of calibrated linear centered kernel alignment (CKA). **b.** Pairwise values of calibrated k-nearest-neighbor (kNN) overlap at *k* = 100. Both panels compare 56 embeddings evaluated on a common set of 14,039 genes. Both matrices show recurring groups of related embeddings, alongside differences in individual pairwise similarities. These differences illustrate the complementary information captured by global gene–gene geometry and local gene neighborhoods.

**Supplementary Figure 2.**
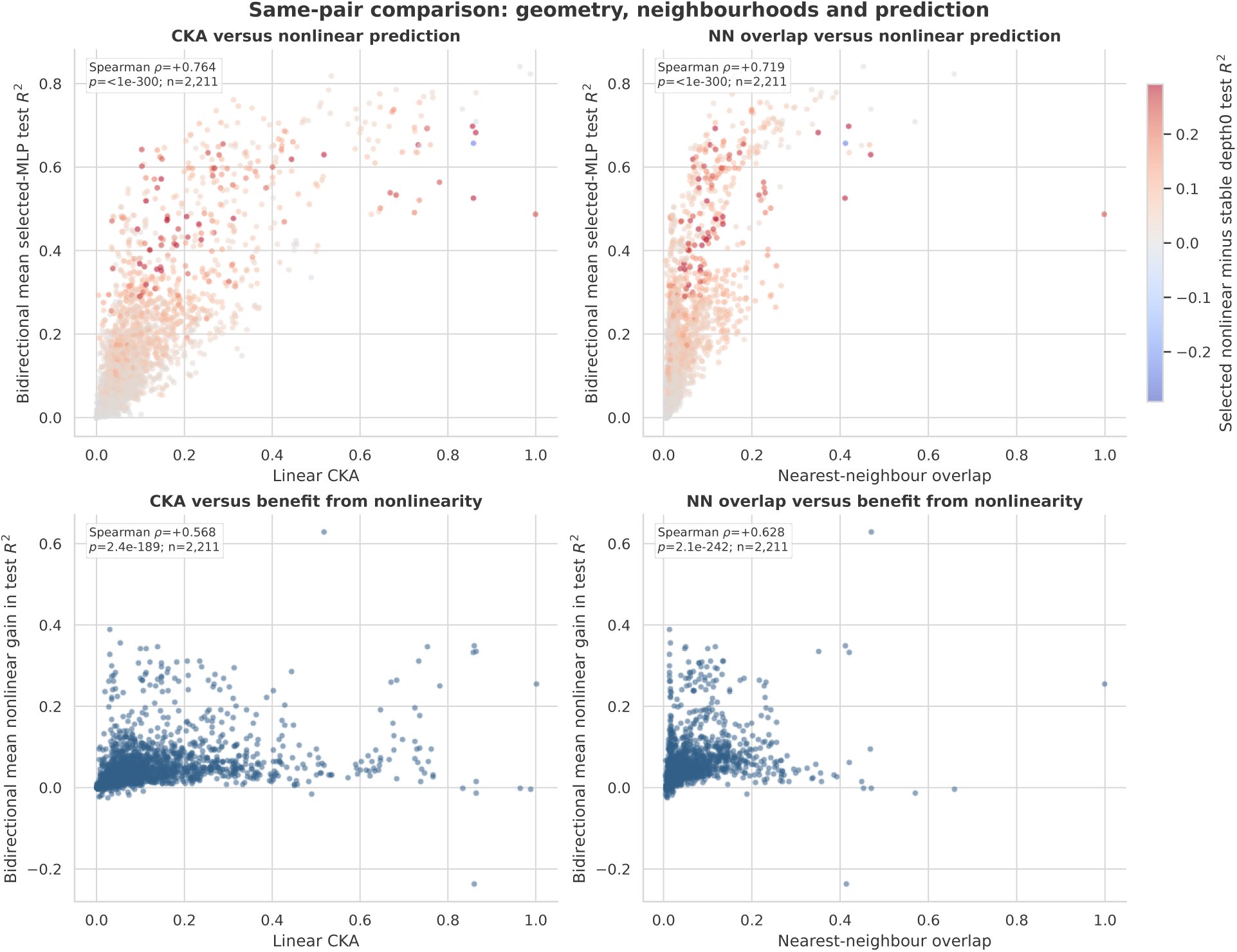
Relationship between embedding concordance and cross-embedding reconstruction. Each point represents one of the 2,211 unordered pairs among the 67 embeddings. The horizontal axes show linear CKA in the left panels and nearest-neighbor overlap in the right panels. In the upper panels, the vertical axis shows the full-target test *R*^2^ of the validation- selected multilayer perceptrons, averaged over the two prediction directions. In the lower panels, the vertical axis shows the corresponding mean gain in test *R*^2^ relative to the linear baseline with no hidden layers. Point colours in the upper panels encode this gain, with red indicating improvement and blue indicating poorer performance than the linear baseline. Embedding pairs with higher global or local concordance generally showed higher bidirectional reconstruction scores. Nonlinear gains also tended to increase with concordance, although their magnitude varied considerably among pairs.

## Supplementary Tables

**Supplementary Table 1.** Characteristics and processing of the 67 core gene embedding sets. For each set, the table reports its modality, number of retained genes, dimensionality, coverage, acquisition route, and processing procedure. Coverage is the percentage of the 20,097 genes in the reference set represented in the final embedding matrix.

| Embeddings | Modality | Genes | Dimensions | Coverage | Acquisition route | Detailed explanation |
|---|---|---|---|---|---|---|
| AAC | Protein sequence | 18,873 | 20 | 93.91% | Downloaded precomputed matrix | Precomputed amino-acid-composition representation. |
| ProtALBERT | Protein sequence | 18,453 | 4,096 | 91.82% | Downloaded precomputed matrix | Precomputed protein language-model representation. |
| AlphaGenome | Regulatory genomics | 20,084 | 256 | 99.94% | Locally generated by model inference | AlphaGenome API regulatory-output tracks for 1,048,576-bp GRCh38 windows centered on strand-aware TSSs. A fixed panel of 250 tracks was summarized over five spatial windows (1,250 features), signed-log1p transformed, feature-wise z-scored, and reduced to 256 components by randomized PCA. These are exposed predictions, not hidden model states. |
| APAAC | Protein sequence | 18,871 | 80 | 93.90% | Downloaded precomputed matrix | Precomputed amphiphilic pseudo-amino-acid-composition representation. |
| Basenji2 | Regulatory genomics | 20,097 | 5,313 | 100.00% | Locally generated by model inference | Basenji2 PyTorch checkpoint basenji2.pth applied to 131-kb GRCh38 sequences centered on each TSS. Human-track predictions were averaged over the central three output bins, log1p transformed, and z-scored feature-wise across genes. |
| ProtBERT-BFD | Protein sequence | 18,453 | 1,024 | 91.82% | Downloaded precomputed matrix | Precomputed protein language-model representation. |
| BERT-PFAM | Protein sequence | 18,461 | 768 | 91.86% | Downloaded precomputed matrix | Precomputed protein language-model representation. |
| BioBERT (gene+protein text) | Biomedical text | 19,168 | 768 | 95.38% | Locally generated by model inference | BioBERT checkpoint dmis-lab/biobert-base-cased-v1.2 applied without fine-tuning to GenePT NCBI and UniProt gene/protein summaries. Final-layer embeddings were mean-pooled over valid non-special tokens (maximum length 512). |
| BioConceptVec CBOW | Biomedical text | 17,040 | 100 | 84.79% | Downloaded precomputed matrix | BioConceptVec CBOW gene-concept vectors from concept_cbow.json. |
| BioConceptVec fastText | Biomedical text | 17,040 | 100 | 84.79% | Downloaded precomputed matrix | BioConceptVec fastText gene-concept vectors from concept_fast.json. |
| BioConceptVec GloVe | Biomedical text | 17,040 | 100 | 84.79% | Downloaded precomputed matrix | BioConceptVec GloVe gene-concept vectors from concept_glove.json. |
| BioConceptVec Skip-Gram | Biomedical text | 17,040 | 100 | 84.79% | Downloaded precomputed matrix | BioConceptVec Skip-gram gene-concept vectors from concept_sg.json. |
| BLAST | Protein sequence | 18,873 | 20,421 | 93.91% | Downloaded precomputed matrix | Precomputed sequence-similarity features. |
| CPC-PROT | Protein sequence | 18,453 | 512 | 91.82% | Downloaded precomputed matrix | Precomputed protein-sequence embedding. It learns fixed-length protein embeddings using contrastive predictive coding . |
| deepNF (STRING) | Networks | 19,071 | 256 | 94.89% | Locally constructed from public data | STRING v12 human evidence-channel networks from the detailed links, protein information, and alias files. Channel-specific sparse PPMI matrices were reduced by TruncatedSVD and combined into a deepNF-inspired 256-dimensional network-fusion representation. |
| DeepSEA | Regulatory genomics | 20,097 | 919 | 100.00% | Locally generated by model inference | DeepSEA through the Kipoi DeepSEA/predict interface applied to 1-kb GRCh38 sequences centered on each TSS. The 919 predicted epigenomic probabilities were z-scored feature-wise across genes. |
| DepMap CRISPR | Functional | 18,562 | 256 | 92.36% | Locally constructed from public data | DepMap CRISPRGeneEffect.csv profiles across cancer models, with Model.csv used for model metadata. The mapped gene-effect matrix was reduced to 256 dimensions by randomized PCA. |
| DNABERT-2 | DNA sequence | 20,085 | 768 | 99.94% | Locally generated by model inference | DNABERT-2 checkpoint zhihan1996/DNABERT-2-117M applied to Ensembl release 114 canonical CDS sequences. Final-layer non-special-token embeddings were mean-pooled; long CDSs were processed in 1,600-nt chunks and combined by sequence-length-weighted averaging. |
| DRKG TransE | Knowledge graph | 18,999 | 400 | 94.54% | Downloaded precomputed matrix | Official 400-dimensional TransE_l2 entity vectors from the DRKG drkg.tar.gz release. |
| Enformer | Regulatory genomics | 20,097 | 5,313 | 100.00% | Locally generated by model inference | TensorFlow Hub Enformer model applied to 393-kb GRCh38 sequences centered on each TSS. Human-head predictions were averaged over the central three output bins, log1p transformed, and z-scored feature-wise across genes. |
| ESM-1b | Protein sequence | 18,453 | 1,280 | 91.82% | Downloaded precomputed matrix | Precomputed protein language-model representation. |
| ESM-2 | Protein sequence | 18,690 | 1,280 | 93.00% | Downloaded precomputed matrix | Precomputed mean-pooled ESM2 protein vectors generated with checkpoint esm2_t33_650M_UR50D and human UniProt protein sequences. |
| FRoGS (ARCHS4-256) | Bulk expression | 16,265 | 256 | 80.93% | Downloaded precomputed matrix | Official FRoGS ARCHS4 co-expression representation from gene_vec_archs4_256.csv. |
| Gene2Vec | Bulk expression | 17,916 | 200 | 89.15% | Downloaded precomputed matrix | Original 200-dimensional Gene2Vec co-expression vectors from gene2vec_dim_200_iter_9.txt. |
| GeneCompass | Multimodal | 16,578 | 768 | 82.49% | Checkpoint parameter extraction | Static learned gene-token vectors extracted from the official GeneCompass Base checkpoint in pretrained_models/GeneCompass_Base, aligned using prior_knowledge/human_mouse_tokens.pickle. No contextual inference, pooling, or fine-tuning was performed. |
| Geneformer | Single-cell expression | 20,007 | 1,152 | 99.55% | Checkpoint parameter extraction | Static gene-token embedding table bert.embeddings.word_embeddings.weight extracted from the official Geneformer V2-316M model.safetensors checkpoint and aligned with token_dictionary_gc104M.pkl. No model inference or pooling was performed. |
| GenePT Ada | Biomedical text | 19,099 | 1,536 | 95.03% | Downloaded precomputed matrix | Precomputed GenePT text-embedding-ada-002 vectors derived from gene summaries. |
| GenePT Model 3 | Biomedical text | 19,103 | 3,072 | 95.05% | Downloaded precomputed matrix | Precomputed GenePT text-embedding-3-large vectors derived from combined NCBI gene and UniProt protein summaries. |
| GO TF-IDF SVD-256 | Functional | 16,833 | 256 | 83.76% | Locally constructed from public data | Direct, non-negated Biological Process annotations from HUMAN-uniprot.gaf.gz. A binary UniProt-by-term matrix was L2 TF-IDF weighted and reduced by randomized TruncatedSVD to 256 dimensions; no ontology-parent propagation was added. |
| GTEx | Bulk expression | 19,353 | 68 | 96.30% | Locally constructed from public data | GTEx v11 median gene-level TPM matrix GTEx_Analysis_2025-08-22_v11_RNASeQCv2.4.3_gene_median_tpm.gct.gz. Tissue features were log1p transformed and z-scored by tissue across mapped genes, with no dimensionality reduction. |
| Helix-mRNA | RNA sequence | 19,976 | 256 | 99.40% | Locally generated by model inference | Helix-mRNA checkpoint helical-ai/helix-mRNA applied to Ensembl release 114 canonical mature mRNA sequences. Real sequence-token embeddings were mean-pooled, and chunk-level vectors were combined using token-count-weighted averaging. |
| Hetionet (non-PPI) | Knowledge graph | 18,787 | 256 | 93.48% | Locally constructed from public data | Hetionet v1.0 edges and nodes from hetionet-v1.0.0.zip, excluding direct PPI context. Typed biomedical neighbors were encoded with TF-IDF, reduced by randomized TruncatedSVD to 256 dimensions, and z-scored across source genes. |
| HMMER | Protein sequence | 18,873 | 20,421 | 93.91% | Downloaded precomputed matrix | Precomputed sequence-profile features. |
| HPA subcellular | Functional | 13,581 | 50 | 67.58% | Locally constructed from public data | Human Protein Atlas subcellular_location.tsv.zip annotations. Reported subcellular locations were encoded as binary multi-hot features and z-scored by location across mapped genes. |
| HPA tissue | Bulk expression | 13,420 | 59 | 66.78% | Locally constructed from public data | Human Protein Atlas normal_ihc_data.tsv.zip annotations. Ordinal protein-expression levels were aggregated by the maximum reported level across cell types within each tissue, then tissue features were z-scored across mapped genes. |
| HyenaDNA | DNA sequence | 20,085 | 256 | 99.94% | Locally generated by model inference | HyenaDNA checkpoint LongSafari/hyenadna-medium-160k-seqlen-hf applied to Ensembl release 114 canonical CDS sequences. Final hidden states were mean-pooled over non-special nucleotide tokens; long CDSs were processed in 12,000-nt chunks and combined by token-count-weighted averaging. |
| Learned-Vec | Protein sequence | 18,873 | 64 | 93.91% | Downloaded precomputed matrix | Precomputed learned-vector representation. |
| Mahi (all-context mean) | Multimodal | 18,918 | 3,666 | 94.13% | Downloaded precomputed matrix | Official precomputed Mahi vectors from mahi_embeddings_float16.parquet. For each uniquely mapped Entrez gene, vectors from the 289 non-global tissue/cell contexts were averaged; the separate global context was excluded. |
| Mahi (global) | Multimodal | 18,918 | 3,666 | 94.13% | Downloaded precomputed matrix | Official precomputed Mahi vectors from mahi_embeddings_float16.parquet. For each uniquely mapped Entrez gene, only the global-context vector was retained; tissue/cell-specific contexts were excluded. |
| Mashup (STRING) | Networks | 16,331 | 800 | 81.26% | Downloaded precomputed matrix | Pretrained Mashup human STRING network vectors and their matching gene list. |
| MSA Transformer (depth 1) | Protein sequence | 19,472 | 768 | 96.89% | Locally generated by model inference | FAIR-ESM MSA Transformer checkpoint esm_msa1b_t12_100M_UR50S applied to the UniProt human reference proteome. Each protein was treated as a one-row MSA and represented by mean-pooled layer-12 residue embeddings. |
| Mut2Vec | Multimodal | 17,445 | 300 | 86.80% | Downloaded precomputed matrix | Precomputed protein-associated representation. |
| NEWT CellNet | Bulk expression | 17,499 | 128 | 87.07% | Downloaded precomputed matrix | Official NEWT CellNet component. |
| NEWT GO graph | Functional | 13,788 | 256 | 68.61% | Downloaded precomputed matrix | Official NEWT GO-graph component. |
| NEWT GO-256 | Functional | 13,788 | 256 | 68.61% | Downloaded precomputed matrix | Official NEWT Gene Ontology component. |
| NEWT MSigDB | Functional | 15,423 | 256 | 76.74% | Downloaded precomputed matrix | Official NEWT pathway/gene-set component. |
| Node2Vec (PPI) | Networks | 18,356 | 128 | 91.34% | Locally constructed from public data | Node2Vec representation constructed from the consensus.dat protein-interaction edge list. Ten unbiased 80-step walks per node (p = q = 1) were used for Word2Vec skip-gram training, producing 128-dimensional vectors with seed 42. |
| Nucleotide Transformer v2 | DNA sequence | 20,085 | 1,024 | 99.94% | Locally generated by model inference | Nucleotide Transformer checkpoint InstaDeepAI/nucleotide-transformer-v2-500m-multi-species applied to Ensembl release 114 canonical CDS sequences. Final-layer embeddings were mean-pooled over valid non-special tokens; 6,000-nt chunks were combined by chunk-aware weighted averaging. |
| OPA2Vec | Functional | 18,660 | 200 | 92.85% | Locally constructed from public data | OPA2Vec-derived 200-dimensional vectors reconstructed in Python 3 from the GO basic ontology and human GOA annotations. Ontology and annotation context was converted to a corpus for Word2Vec-style training. |
| Orthrus | RNA sequence | 19,976 | 256 | 99.40% | Locally generated by model inference | Orthrus checkpoint quietflamingo/orthrus-base-4-track applied to Ensembl release 114 canonical mature transcripts. Full-sequence representations were used when possible; longer transcripts were chunked and combined by length-weighted averaging. |
| PrimeKG (non-PPI) | Knowledge graph | 19,216 | 256 | 95.62% | Locally constructed from public data | PrimeKG kg.csv edges, excluding direct PPI context. Typed biomedical neighbors were encoded with TF-IDF, reduced by randomized TruncatedSVD to 256 dimensions, and z-scored across source genes. |
| ProtVec | Protein sequence | 18,873 | 100 | 93.91% | Downloaded precomputed matrix | Precomputed ProtVec protein representation. |
| PubMedBERT (gene+protein text) | Biomedical text | 19,168 | 768 | 95.38% | Locally generated by model inference | PubMedBERT/BiomedBERT checkpoint microsoft/BiomedNLP-BiomedBERT-base-uncased-abstract-fulltext applied without fine-tuning to GenePT NCBI and UniProt gene/protein summaries. Final-layer embeddings were mean-pooled over valid non-special tokens (maximum length 512). |
| RNA-FM | RNA sequence | 19,976 | 640 | 99.40% | Locally generated by model inference | RNA-FM checkpoint RNA-FM_pretrained.pth applied to Ensembl release 114 canonical mature RNA sequences. Layer-12 embeddings were mean-pooled excluding BOS/EOS tokens, and long-transcript chunk vectors were combined by sequence-length-weighted averaging. |
| scFoundation | Single-cell expression | 19,062 | 768 | 94.85% | Checkpoint parameter extraction | Fixed gene-position/identity table model.pos_emb.weight extracted from the official scFoundation models.ckpt checkpoint and aligned with OS_scRNA_gene_index.19264.tsv. No cell-context inference or pooling was performed. |
| scGPT (PanCancer) | Single-cell expression | 19,071 | 512 | 94.89% | Checkpoint parameter extraction | Static gene-token table encoder.embedding.weight extracted from the official scGPT pan-cancer best_model.pt checkpoint and aligned with vocab.json. No contextual inference, pooling, or fine-tuning was performed. |
| scGPT (Whole) | Single-cell expression | 19,280 | 512 | 95.93% | Checkpoint parameter extraction | Static gene-token table encoder.embedding.weight extracted from the official scGPT whole-human best_model.pt checkpoint and aligned with vocab.json. No contextual inference, pooling, or fine-tuning was performed. |
| scPRINT | Single-cell expression | 19,995 | 256 | 99.49% | Checkpoint parameter extraction | Static gene-token table gene_encoder.embedding.weight extracted from the official medium-v1.5.ckpt scPRINT checkpoint and aligned with the checkpoint’s embedded gene list. No contextual or denoised model output was used. |
| SeqVec | Protein sequence | 18,873 | 1,024 | 93.91% | Downloaded precomputed matrix | Precomputed SeqVec protein representation. |
| SpliceAI | Regulatory genomics | 20,097 | 606 | 100.00% | Locally generated by model inference | SpliceAI applied to GRCh38 sequence using GENCODE v49 exon coordinates and canonical transcript identifiers. Canonical splice-site predictions were aggregated into one fixed 606-feature profile per gene and z-scored feature-wise across genes. |
| STRING GNN | Networks | 18,028 | 256 | 89.70% | Downloaded precomputed matrix | Precomputed 256-dimensional residual-GCN node embeddings learned by GenBio AI from a filtered STRING v12.0 network with a link-prediction objective. |
| STRING WaveGC | Networks | 18,131 | 128 | 90.22% | Downloaded precomputed matrix | Precomputed 128-dimensional WaveGC node embeddings learned from the STRING network with a link-prediction objective. |
| ProtT5 | Protein sequence | 18,453 | 1,024 | 91.82% | Downloaded precomputed matrix | Precomputed protein representation. |
| TCGA | Bulk expression | 18,221 | 50 | 90.67% | Downloaded precomputed matrix | Precomputed cancer-expression representation. |
| UCE-33L | Multimodal | 19,082 | 1,280 | 94.95% | Checkpoint parameter extraction | Fixed 1,280-dimensional UCE gene-token vectors generated from the official human segment of all_tokens.torch and the UCE encoder projection. Contextual transformer layers were not used. |
| UniRep | Protein sequence | 18,873 | 5,700 | 93.91% | Downloaded precomputed matrix | Precomputed UniRep protein representation. |
| ProtXLNet | Protein sequence | 18,453 | 1,024 | 91.82% | Downloaded precomputed matrix | Precomputed protein representation. |

**Supplementary Table 2.** Identifier harmonization outcomes for precomputed embeddings.

| **Embedding** | **Uniquely mapped rows** | **Ambiguous rows** | **Dropped rows** | **Merged rows** |
|---|---|---|---|---|
| AAC | 18,905 | 186 | 1,516 | 32 |
| ProtALBERT | 18,482 | 186 | 1,514 | 29 |
| APAAC | 18,903 | 186 | 1,391 | 32 |
| ProtBERT-BFD | 18,482 | 186 | 1,514 | 29 |
| BERT-PFAM | 18,490 | 186 | 1,517 | 29 |
| BioConceptVec CBOW | 17,042 | 32 | 127,542 | 2 |
| BioConceptVec fastText | 17,042 | 32 | 127,542 | 2 |
| BioConceptVec GloVe | 17,042 | 32 | 127,542 | 2 |
| BioConceptVec Skip-Gram | 17,042 | 32 | 127,542 | 2 |
| BLAST | 18,905 | 186 | 1,516 | 32 |
| CPC-PROT | 18,482 | 186 | 1,514 | 29 |
| DRKG TransE | 19,002 | 36 | 20,218 | 3 |
| ESM-1b | 18,482 | 186 | 1,514 | 29 |
| ESM-2 | 18,708 | 160 | 195 | 18 |
| FRoGS (ARCHS4-256) | 16,271 | 148 | 6,699 | 6 |
| Gene2Vec | 17,965 | 32 | 6,482 | 49 |
| GenePT Ada | 19,208 | 1,183 | 74,592 | 109 |
| GenePT Model 3 | 19,583 | 2,088 | 114,153 | 480 |
| HMMER | 18,905 | 186 | 1,516 | 32 |
| Learned-Vec | 18,905 | 186 | 1,516 | 32 |
| Mahi (all-context mean) | 18,918 | 155 | 155 | 0 |
| Mahi (global) | 18,918 | 155 | 155 | 0 |
| Mashup (STRING) | 16,359 | 23 | 2,003 | 28 |
| Mut2Vec | 17,446 | 93 | 1,138 | 1 |
| NEWT CellNet | 17,506 | 121 | 4,150 | 7 |
| NEWT GO graph | 13,795 | 88 | 587 | 7 |
| NEWT GO-256 | 13,795 | 88 | 587 | 7 |
| NEWT MSigDB | 15,427 | 69 | 4,081 | 4 |
| ProtVec | 18,905 | 186 | 1,516 | 32 |
| SeqVec | 18,905 | 186 | 1,516 | 32 |
| STRING GNN | 18,028 | 0 | 838 | 0 |
| STRING WaveGC | 18,131 | 0 | 98 | 0 |
| ProtT5 | 18,482 | 186 | 1,514 | 29 |
| TCGA | 18,251 | 157 | 695 | 30 |
| UniRep | 18,905 | 186 | 1,516 | 32 |
| ProtXLNet | 18,482 | 186 | 1,514 | 29 |

**Supplementary Table 3.** Definitions of the considered modality categories. The 11 categories describe the principal biological information used to construct each representation.

| **Modality** | **Definition** |
|---|---|
| Biomedical text | Embeddings derived from biomedical literature, gene or protein descriptions, or other textual corpora using language models or word-embedding methods. |
| Bulk expression | Embeddings derived from RNA or protein expression profiles across tissues or bulk samples, including those based on co-expression relationships. |
| DNA sequence | Embeddings derived from gene-associated DNA sequences primarily generated from coding DNA sequences. |
| Functional | Embeddings derived primarily from functional annotations, curated gene sets, subcellular localization, or functional assays such as CRISPR dependency screens. |
| Knowledge Graph | Embeddings derived from heterogeneous biomedical graphs containing multiple entity and relationship types, such as genes, diseases, drugs, pathways, and phenotypes. |
| Multimodal | Embeddings constructed or learned by combining multiple types of biological information. |
| Networks | Embeddings derived primarily from the topology of gene or protein interaction networks, particularly protein–protein interaction networks. |
| Protein sequence | Embeddings derived primarily from amino-acid sequences, including embeddings produced by protein language models, sequence-alignment and similarity methods, protein-family models, and amino-acid composition descriptors. |
| RNA sequence | Embeddings derived directly from full-transcript or messenger-RNA sequences using RNA sequence models. |
| Regulatory genomics | Embeddings summarizing regulatory or splicing information obtained from gene-associated genomic regions, including predicted transcriptional, epigenomic, and splice-site features. |
| Single-cell expression | Gene-token embeddings learned by pretrained models from large collections of single-cell transcriptomic profiles. |

